# Initiation of ensemble kinesin-3 motility is regulated by the rigidity of cargo-motor attachment

**DOI:** 10.1101/2021.12.02.470904

**Authors:** Prakash Lama, Minhajuddin Sirajuddin

**Affiliations:** Centre for Cardiovascular Biology and Disease, Institute for Stem Cell Science and Regenerative Medicine, GKVK Campus, Bengaluru – 560065, India; Manipal Academy of Higher Education, Manipal, India

**Author notes:** Address correspondence to M. Sirajuddin.

## Abstract

Intracellular cargo transport is powered by molecular motors that move on their respective filamentous tracks. A key component in this process is the tether between cargo and motor, which is often connected by long slender coiled-coils. Several studies have identified mechanisms that regulate cargo transport and can be broadly categorized into regulation of the motor ATPase activity by autoinhibition, cargo adapters and modifications in the cytoskeletal tracks. The regulatory effects of cargo-motor linkers have been described in kinesin-3 subfamily motors. However, the effects of cargo-motor linker rigidity on ensemble cargo transport has not been explored. Here we have built a DNA origami scaffold, which can be tethered with multiple kinesin-3 motors using either single or double-stranded DNA linkages, mimicking rigid versus flexible cargo-motor linkages. Using this system, we show that regardless of the motor numbers attached to the cargo, only linkers with a lesser degree of freedom allow motors to engage with microtubule tracks. Together, our work identifies that the rigidity of cargo-motor linkages influences motor motility. This opens up the possibilities to identify new factors that can influence the rigidity of cargo-motor linkages that in turn can regulate intracellular cargo transport.

## Introduction

Molecular motors belonging to the kinesin superfamily require three important components for their cellular function; the motor domain, coiled-coil helices and the terminal cargo binding domain (Sweeney and Holzbaur, 2018). The kinesin motor domain interacts with the microtubules, which in turn stimulates ATP turnover and the ATP hydrolyzing activity is converted into the mechanical force required for motility. Modulation of ATPase activity (Gennerich and Vale, 2009), sequestration of the motor domain (Hammond et al., 2010; Imanishi et al., 2006; Toropova et al., 2017; Ren et al., 2018) and modifications in the microtubule tracks (Sirajuddin et al., 2014; Lessard et al., 2019; McKenney et al., 2016; Monroy et al., 2020) are well-known regulatory mechanisms that govern kinesin motility. The cargo binding domain of a typical kinesin motor is located at the distal end of the motor domain. In many cases, the sequestration of motor domain is mediated by the cargo binding domain leading to an autoinhibitory state. This autoinhibition can be relieved by cognate cargo adapters, thereby ensuring fidelity and minimizing futile ATPase cycles during intracellular transport (Siddiqui and Straube, 2017). The third component in kinesin molecular motors are the tethers that connect motor domain and cargo binding domain. These tethers are often dimeric coiled-coil helices that span several hundreds of amino acids, thus rendering a single kinesin molecular motor dimeric in nature (Hirokawa et al., 2009b). The slender coiled coil domain also contains several breaks making them flexible, thereby providing several degrees of freedom to the kinesin motor during cargo transport.

Among the kinesin superfamily of motors, members of the kinesin-3 subfamily such as KIF1A, KIF13B and KIF16 are known to be regulated by the flexible parts of the coiled coil domain (Soppina et al., 2014). Using single molecule and engineered kinesin-3 motors it was inferred that the flexibility of the hinge region between NC and CC1 domains is important for dimerization and subsequent motility of kinesin-3 motors (Soppina et al., 2014; Huo et al., 2012). The dimerization requirement for kinesin-3 processive motility was a contentious notion (Okada and Hirokawa, 1999; Hirokawa et al., 2009a), which has been sufficiently addressed by several independent studies (Soppina et al., 2014; Hammond et al., 2009; Tomishige et al., 2002). The current model suggests that dimerization of kinesin-3 motors leads to super-processive motility i.e., their ability to walk long distances along the microtubules (Soppina et al., 2014; Scarabelli et al., 2015; Siddiqui and Straube, 2017). The super-processive property of kinesin-3 motors have been attributed to the lysine rich loop (K-loop) in the motor domain (Soppina and Verhey, 2014). The basic charge of K-loop mediates electrostatic interaction with the acidic carboxy-terminal tails of tubulin, thus increasing the processive motility of kinesin-3 motors. A recent study has also shown that multimerization of kinesin-3 monomers can also result in cargo transport (Schimert et al., 2019). While tremendous progress has been achieved in understanding kinesin-3 motors and their motility at the single-molecule level, the regulatory aspects of multiple kinesin-3 motors have not been explored. In fact, the conventional method to study molecular motors and their regulation has been an endeavor that majorly involves purified molecular motors and studying them at the single-molecule level. However, molecular motors often work in teams with varying motor ensemble numbers. To study motor ensemble properties and probe the importance of the cargo-motor linker rigidity in ensemble conditions we designed a synthetic scaffold system. The DNA origami scaffold developed in this study can accommodate up to 28 motors, far exceeding the ensemble numbers achieved in previous studies (Toropova et al., 2017; Derr et al., 2012; Driller-Colangelo et al., 2016; Hariadi et al., 2015b; Furuta et al., 2013). Using this system, we probed the role of motor-cargo linkers and found that the initiation of kinesin-3 ensemble motility is dictated by the flexibility of the linkers. Thus, the DNA origami tool described here becomes a robust system that can be extended to study regulatory mechanisms of other molecular motors in their ensemble state.

## Results

### Design and validation of 6HB-400nm DNA origami scaffold

Molecular motors have been extensively studied at the single molecule level, however while performing their cellular functions they often work in ensembles. Few studies have created synthetic DNA scaffolds to study motor ensembles and so far, the systems that exist can assemble maximum of 6 motors with regular spacing (Derr et al., 2012; Hariadi et al., 2015b). To expand the capabilities in studying motor ensembles, here we adapted a DNA origami scaffold that was designed earlier (Bui et al., 2010). The DNA origami scaffold used in this study is 400nm long with 6 helix bundles (6HB-400nm) (Methods) (Figure 1A & Supplementary Figure 1). The 6HB-400nm scaffold has been designed to accommodate up to 28 oligonucleotide-overhangs (called handles) in a linear stretch, each handle is spaced ∼14nm apart (Supplementary Figure 1). The 6HB-400nm scaffold was validated to confirm its structure and occupancy of oligonucleotide handles using anti-handles conjugated with fluorescent molecules and motors at helix 1 & 5 and helix 2 respectively (Methods) (Supplementary Table 1 & Supplementary Figure 2A-D). For the protein attachment to the 6HB-400nm scaffold we designed two versions; A 40 and 20 base single-stranded oligonucleotide (anti-handle) that is covalently linked to the tail end of kinesin through SNAP-tag (Methods) (Supplementary Figure 3A & 3B). The 20 base oligonucleotides anti-handle is fully complementary to the handle sites emanating from the 6HB-400nm scaffold mimicking the rigid linker (Supplementary Figure 3C). In the case of 40 base oligonucleotides anti-handle, the complementarity is restricted only towards 20 basepairs at the 5` of the handle sites rendering them as flexible linkers. On the other hand, the handle oligonucleotides that are complementary to the anti-handles are also 40 base long of which 20 base towards the 3`/5` end (i.e., towards the DNA scaffold) remain single-stranded (Supplementary Figure 3C). From hereafter we refer the 20 and 40 oligonucleotides protein-DNA scaffold attachment linkers as rigid and flexible linkers respectively (Supplementary Figure 3C).

**Figure 1:**
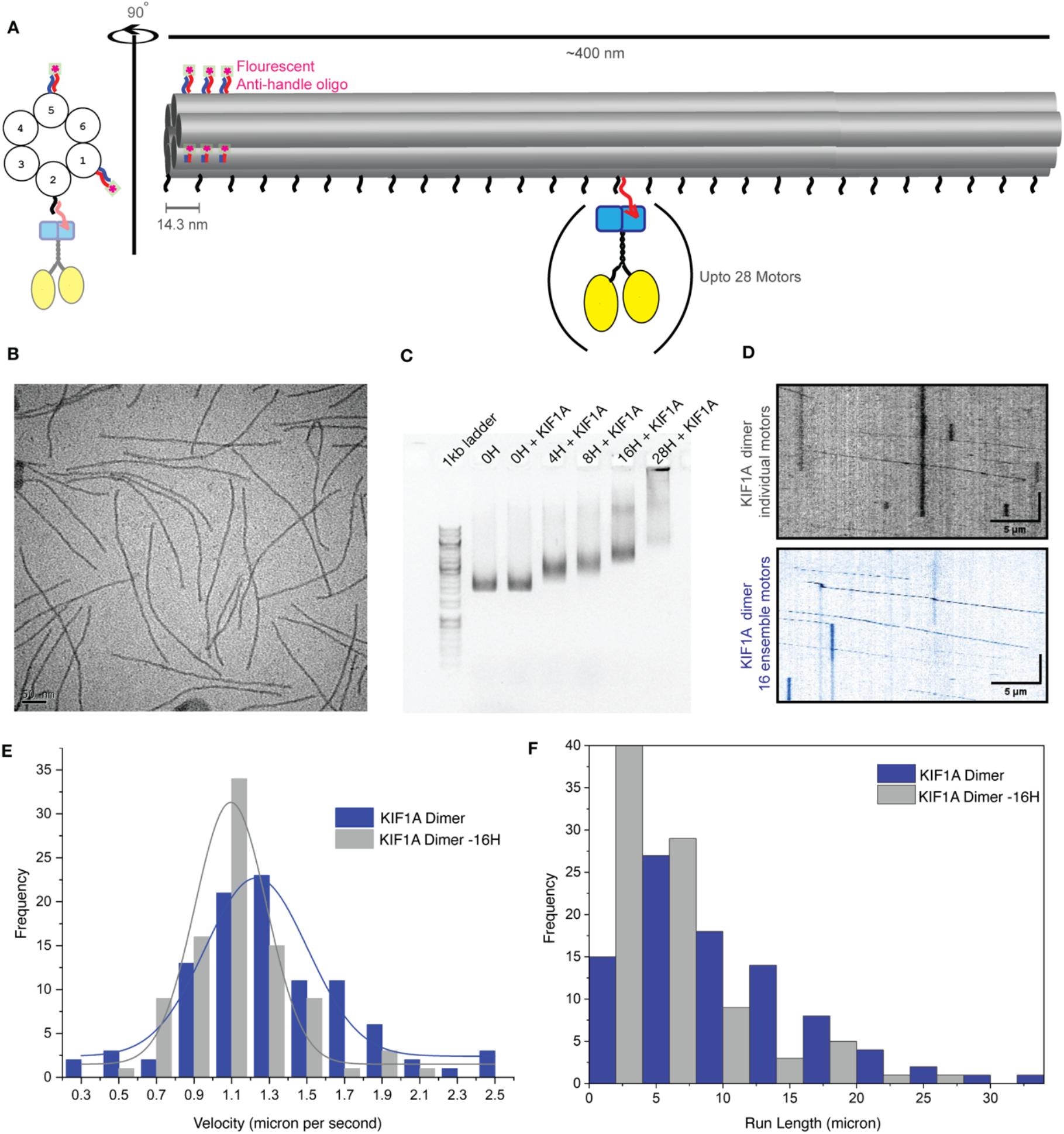
Characterization of 400nm-6HB DNA origami as motor cargo scaffold. **A**. Cartoon representation of 6HB-400nm DNA origami scaffold used in this study as illustrated. 28 handle sites are located in the helix 2 as a single file and three fluorescent handles are located each in helix 1 and 5. **B**. Negative stained electron micrograph of 6HB-400nm DNA origami scaffold. Scale bar = 50nm. **C**. Agarose gel shift assays for 6HB-400nm scaffold with and without KIF1A dimers for varying handle sites as marked. **D**. Representative kymographs of KIF1A 1-393 SNAP-647 (grey) and 6HB-400nm 28H KIF1A 1-393 SNAP ensemble motors (blue) moving on microtubules. Scale bar = 5μm and 10 seconds. **E**. Velocity and processivity histograms of KIF1A 1-393 SNAP-647 (in grey) and 6HB-400nm 16H KIF1A 1-393 SNAP ensemble (in blue) motors. The average velocity of KIF1A 1-393 SNAP-647 are 1.27±0.39μm s^-1^, n=100 6HB-400nm 16H KIF1A 1-393 SNAP ensembles are 1.08±0.29μm s^-1^, n=91. The average run length of KIF1A 1-393 SNAP-647 are 3.36μm and 6HB-400nm 16H KIF1A 1-393 SNAP ensembles are 9.8μm. n represents the number of motor particles analyzed.

Next, we established motor motility assays assessing the feasibility of the 6HB-400nm scaffold to study the ensemble motor behavior. Briefly, KIF1A motors were covalently linked to single-stranded DNA oligonucleotides (flexible and rigid anti-handles) that are complementary to the handles present in the 6HB-400nm scaffold (Methods) (Figure 1C and Supplementary Figure 3A & 3B). To visualize the 6HB-400nm scaffolds in our motility assays, we labelled them using six fluorescent oligonucleotides that were attached to the fluorescent handles at helix 1 & 5 (Methods) (Figure 1A, Supplementary Figure 1 & Supplementary Table 1). In our motility assays, we compared the 6HB-400nm:28 KIF1A motors (ensemble motors) motility against individual KIF1A motors (Methods). Our results show that the velocity of 6HB-400nm:28 KIF1A ensemble and single KIF1A motors are 1.27±0.39 μm s^-1^ and 1.08±0.29 μm s^-1^ respectively (Figure 1D & 1E) (Supplementary Table 2). In the case of processivity, the 6HB-400nm:16 KIF1A ensembles and single KIF1A motors are near identical (Figure 1F) (Supplementary Table 1). From this experiment we conclude that the 6HB-400nm scaffold can serve as a tool for characterizing motor ensemble properties.

### Motility properties of KIF1A motor ensembles

Full-length Kinesin-3 motors can exist as monomeric units and homodimers in cells, single-molecule studies have suggested that the dimeric KIF1A motors are super-processive (Soppina et al., 2014; Tomishige et al., 2002). However, a quantitative comparison of monomer versus dimer KIFA ensembles is yet to be characterized. Moreover, the flexibility of cargo-motor linkers in an ensemble setting has not been explored. In order to test this, we compared the ensemble motility properties of dimeric and monomeric KIF1A tethered to 6HB-400nm scaffold, hereafter called as 6HB-400nm monomer KIFA and dimer KIF1A ensembles respectively (Methods) (Supplementary Figure 3A). We chose four different ensemble numbers in this study; 28, 16, 8 and 4 motor attachment handle sites, which we hereafter refer to as 28H, 16H, 8H and 4H respectively (Methods). The 6HB-400nm:KIF1A dimer ensembles with varying handle sites and linkers were analyzed in an agarose gel to confirm the extent of DNA:motor complex formation (Methods) (Figure 1C, Supplementary Figure 3E & 3F).

From the motility assays, we could record individual 6HB-400nm:KIF1A ensembles moving along the microtubules, regardless of the KIF1A oligomeric state, ensemble number and the nature of linkers. Upon quantification of velocity and processivity values, deviations between different 6HB-400nm:KIF1A ensembles began to emerge (Figure 2 & Supplementary Figure 4 & 5). The velocities within monomer and dimer KIF1A ensemble cohorts does not change with motor number. However, the velocity between monomer and dimer KIF1A rigid ensembles are two-fold different; for 6HB-400nm: 4-28H dimer and monomer KIF1A rigid ensembles the average velocity is ∼2 μm s^-1^ and ∼0.9 μm s^-1^ respectively (Figure 2A, Supplementary Figure 4 & Supplementary Table 2). Compared to the KIF1A monomer rigid ensembles the flexible cohort exhibited two-fold reduction in velocity (Figure 2A, Supplementary Figure 4 & Supplementary Table 2). In contrast, the processivity values follow similar trend between monomer rigid, monomer flexible and dimer rigid ensembles (Figure 2B, Supplementary Figure 4 & Supplementary Table 2). A puzzling phenomenon here is the poor processivity of KIF1A ensembles below 16H. The 6HB-400nm-4H and 8H KIF1A dimer processivity is 4.96±0.01 μm and 4.95±0.01 μm respectively. We attribute this low processivity of 6HB-400nm-4H & 8H KIF1A dimer ensembles to the high magnesium concentration used in our motility assays. High magnesium levels are required to overcome the magnesium quenching property of 6HB-400nm DNA scaffolds and therefore balancing the requirement of Mg2+ ions for motor ATPase activity (Methods). Indeed, when high magnesium is used in motility assays the single dimeric KIF1A also exhibits poor processivity of 3.46±0.01 μm (Figure 1E & Supplementary Table 2). We also observed that the high magnesium induced low processivity can be overcome by higher motor ensembles. For example, the processivity of 6HB-400nm dimer KIF1A rigid 16H and 28H ensembles are 12.2±0.01 μm and 7.2±0.01 μm respectively and is similar to the dimeric KIF1A processivity 9.8±0.01 μm (Supplementary Table 2).

**Figure 2:**
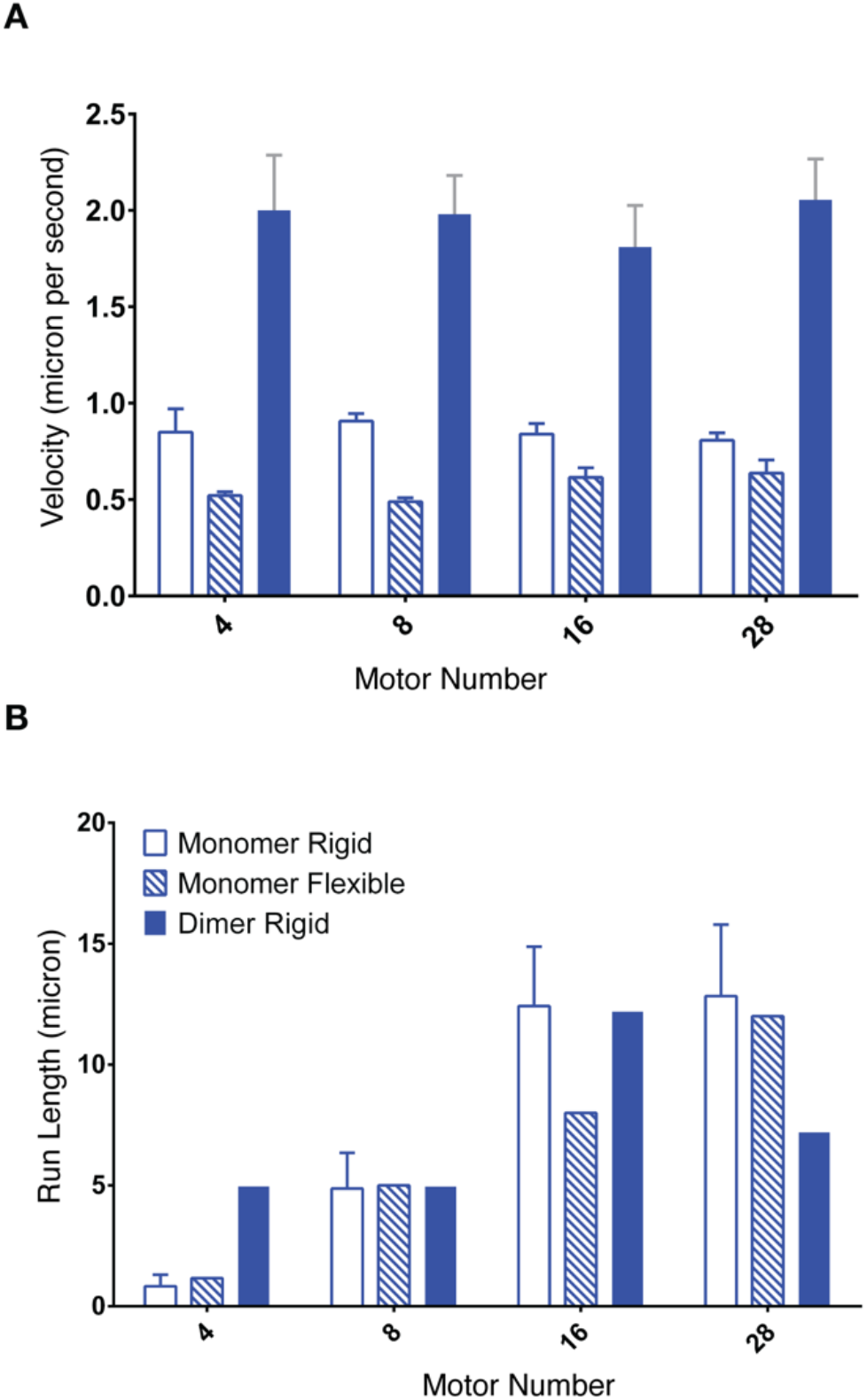
KIF1A dimer versus monomer ensemble motility. **A**. Average velocity data of 6HB-400nm 4-28H KIF1A monomer ensembles with flexible and rigid oligonucleotides and KIF1A dimer ensembles with rigid oligonucleotides. **B**. Average processivity values of 6HB-400nm 4-28H KIF1A ensembles as described in A. Error bars represent the standard error of the mean from three independent experiments. For individual values see Supplementary Table 2.

In summary, the velocity of KIF1A ensembles remains unaffected regardless of the motor number and the processivity of KIF1A ensembles improves as the motor number increases.

### Comparative analysis of flexible versus rigid cargo-motor linkers

While the velocity and processivity of flexible versus rigid cargo-motor linkers in ensembles remain largely unchanged, we observed reduced number of 6HB-400nm KIF1A monomer ensembles with flexible linkers encountering the microtubule (Supplementary Movie 1). To gain more insights, we qualitatively assessed the ability of flexible and rigid linked 6HB-400nm KIF1A monomer ensembles binding microtubules in the absence of ATP, a microtubule strong binding condition called rigor-state (Methods). The rigor-state assays exemplify our observation that the 6HB-400nm KIF1A monomer ensembles with flexible linkers show reduced binding to the microtubules (Figure 3). In order to systematically probe the cargo-motor linkers, we additionally generated linkers that have single-stranded stretch at the either end of scaffold or motor tail, called intermediate flexible linkers (Methods) (Supplementary Table 1). In the rigor-state assays the 6HB-400nm KIF1A monomer ensembles linked with intermediate flexible linkers also showed a marked decrease in microtubule binding (Figure 3). The reduced binding of 6HB-400nm KIF1A monomer ensembles is a recurring phenomenon regardless of motor number sampled in our assays (Figure 3 & Supplementary Figure 6).

**Figure 3:**
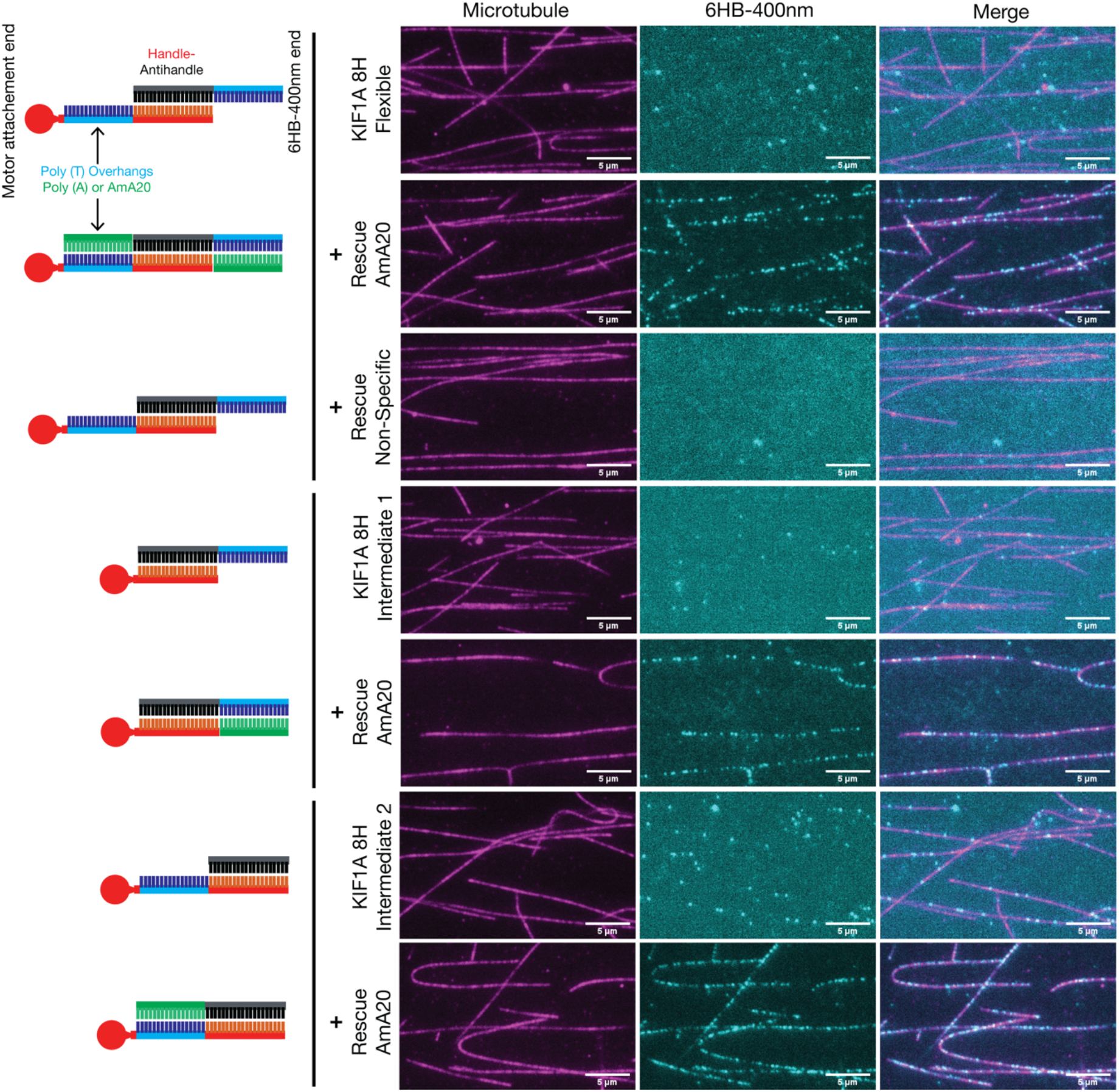
Microtubule-binding of KIF1A ensembles with varying linkers. Microtubule-binding of 6HB-400nm 8H KIF1A monomer ensembles with varying linkers as indicated and illustrated. The microtubules are shown in magenta and the 6HB-400nm KIF1A 8H in cyan. In each microtubule-binding experiment, a rescue reaction was perfumed (marked as rescue), where an oligonucleotide that is complementary to the flexible parts of linkers was added to the mixture. For more details regarding the sequences of linkers and rescue oligonucleotides see Supplementary Table 1. Scale bar = 5μm.

Next, we reasoned whether the reduced microtubule-binding ability of 6HB-400nm KIF1A monomer ensembles with flexible linkers could be rescued. For which we designed rescue oligonucleotides that can basepair with the single-stranded stretches in the flexible linkers (Methods) (Figure 3, Supplementary Figure 6 & Supplementary Table 1). The rigor-state assays were performed in the absence and presence of rescue oligonucleotides for flexible 6HB-400nm monomer KIF1A ensembles (Methods). Remarkably, the microtubule-binding of 6HB-400nm KIF1A monomer ensembles with the flexible linkers was restored upon the addition of rescue oligonucleotides (Figure 3). The microtubule-binding rescue was observed with 16H also, however, the effects were more pronounced with 8H (Figure 3 & Supplementary Figure 6). To establish whether the rescue is specific, we performed the rigor-state rescue experiments in the presence of a scrambled oligonucleotides (Supplementary Table 1). In comparison to the rescue oligonucleotides that have complementarity to the flexible linkers the scrambled oligonucleotides did not show any improvement in microtubule-binding of 6HB-400nm KIF1A monomer ensembles (Figure 3 & Supplementary Figure 6). These experiments conclusively demonstrate that the microtubule-binding of KIF1A monomer ensembles can be regulated by the stiffness of the cargo-motor linker.

### Initiation of KIF1A ensemble motility is sensitive to the rigidity of DNA linkers

The above rescue experiments were performed in the rigor-state. To assess the functional significance of cargo-motor linker flexibility, we next performed experiments in the presence of ATP (Methods). In addition to the velocity and processivity values, we measured the binding frequency of flexible 6HB-400nm KIF1A monomer ensembles with and without the rescue oligonucleotides (Methods). The velocity and processivity values of flexible 6HB-400nm KIF1A monomer ensembles do not change upon addition of rescue oligonucleotides (Figure 4A, Supplementary Figure 7 & Supplementary Table 1), similar to our results with flexible versus rigid linkers (Figure 2). However, we observed a marked decrease in landing frequency with the flexible 6HB-400nm KIF1A monomer ensembles (Figure 4B). The landing frequency rate of flexible 6HB-400nm 4H, 8H and 16H KIF1A ensembles are 0.004, 0.001 and 0.007 μm^-1^ s^-1^ respectively. Similar to the rigor-state experiments, we performed the motility experiments with flexible 6HB-400nm KIF1A monomer ensembles in the presence of rescue oligonucleotides (Methods). The number of flexible 6HB-400nm KIF1A monomer ensemble particles moving on the microtubules markedly increases with rescue oligonucleotides (Supplementary Movie 1). Upon quantification, the landing-frequency of flexible 6HB-400nm KIF1A monomer ensembles significantly increases with the addition of rescue oligonucleotides in the motility assays (Figure 4B & Supplementary Figure 7). The landing frequencies of rescued 6HB-400nm 4H, 8H and 16H KIF1A monomer ensembles are 0.011, 0.073 and 0.059 μm^-1^ s^-1^ respectively. We also observed the improvement in landing frequency is specific to the rescue oligonucleotides that are complimentary to the single-stranded DNA linkers. Combinedly these experiments suggest that the initiation of KIF1A ensembles motility is sensitive to the rigidity of the linkers between the 6HB400nm scaffold and motors.

**Figure 4:**
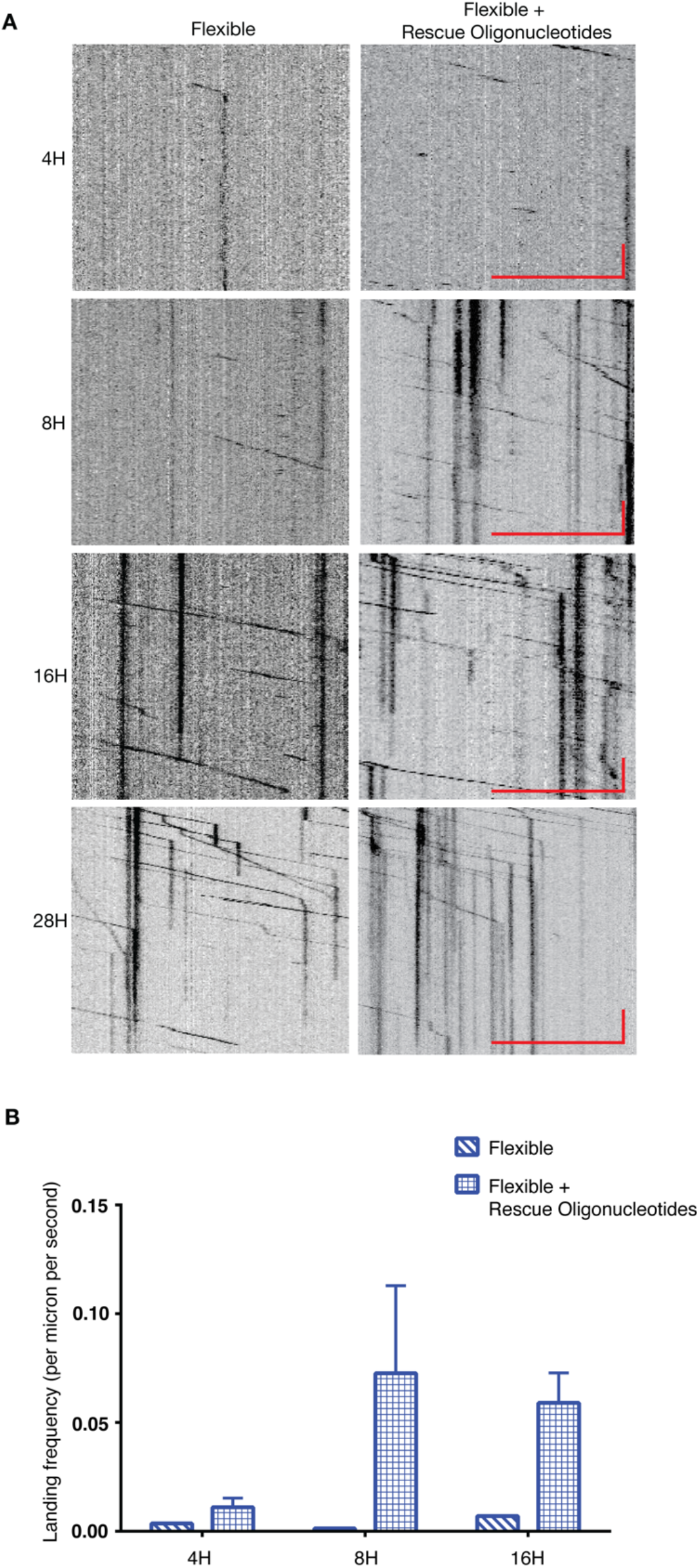
Motility properties of KIF1A ensembles with varying linkers. **A**. Representative kymographs of 6HB-400nm 4-28H KIF1A monomer ensembles with flexible and flexible-rescue oligonucleotides. Scale bar = 5μm and 10 seconds. **B**. Average landing frequency rates for 6HB-400nm 4H, 8H and 16H KIF1A monomer ensembles with flexible linkers and rescue oligonucleotides as indicated. Error bars represent the standard error of the mean from three independent experiments.

## Discussion

Majority of the motor regulation studies have been performed using purified components of single or individual motor proteins. There are a few exceptions where the tug-of-war between two opposing motors have been described (Derr et al., 2012; Hariadi et al., 2015b; Toropova et al., 2017; Driller-Colangelo et al., 2016; Furuta et al., 2013). However, the regulatory aspects of ensemble motors have been poorly described, for instance the effects of cargo-motor linker flexibility towards ensemble kinesin-3 motility is unknown. Similarly, the ensemble motor experiments so far has been limited to DNA origami scaffolds with six motors. On the other hand, DNA scaffolds that can accommodate hundreds of motors has been described. However, it can only control the spacing between each motor thus limiting the ability to control the motor ensemble numbers (Hariadi et al., 2015a). Molecular motors that work together have their ensemble number in varying orders. For example, the motor ensemble numbers involved in the intracellular and intraflagellar cargo transport can range from 2 – 20 (Hirokawa et al., 2009b; Rai et al., 2016; Siddiqui and Straube, 2017; Prevo et al., 2017), and in the sarcomere more than 50 muscle myosins can engage during muscle contraction (Spudich, 2014). In order to expand the capabilities to understand the molecular motor ensembles in this study we have designed and validated a 6HB-400nm DNA origami scaffold. The 6HB-400nm DNA origami scaffold also acts as a cargo mimic, where the cargo-motor linkers are amenable for varying degrees of flexibility. Therefore, the 6HB-400nm DNA origami scaffold described here offers versatility to study molecular motor ensembles up to 28 molecules.

Using 6HB-400nm DNA origami scaffold as a cargo, we tethered them with the KIF1A motors with varying ensemble motor numbers and linker flexibilities. From our assays we show that the biochemical properties of the KIF1A motors does not get affected as evident from the velocity and processivity values between rigid and flexible linkers. However, a striking observation emerged from these assays is the inability of KIF1A monomer ensembles to initiate motility when tethered to 6HB-400nm with flexible linkers (Figure 3 and 4). We also unequivocally show that the effects of flexible linkers can be reversed by the addition of oligonucleotides that are complementary to the flexible single-strand DNA linkers i.e., the flexible linkers (Figure 3 and 4). We attribute the mechanism of diminished microtubule-binding of KIF1A ensembles with flexible linkers to its autoinhibited state mediated by the K-loop. In the case where the 6HB-400nm KIF1A monomer ensembles tethered with flexible oligonucleotides, the motor domain can adopt several degrees of conformation with respect to the 6HB-400nm scaffold. This might be conducive for an electrostatic interaction between the negatively charged DNA strands of 6HB-400nm and the positively charged K-loop of KIF1A motor domain. Upon addition of the rescue oligonucleotides complementary to the flexible regions the persistence of the linkers decreases. Thus, the motor domain may remain with a restricted degree of conformational flexibility and no electrostatic interaction between K-loop and 6HB-400nm DNA elements.

The K-loop of kineins-3 motors is a unique element that has been shown to enhance the processivity among this subfamily of kinesin motors (Soppina and Verhey, 2014). This is mediated by the electrostatic interaction with the acidic carboxy-terminal tails of alpha- and beta-tubulins. Therefore, it is conceivable that such an electrostatic interaction between K-loop and other acidic elements might sterically interfere with microtubule interaction. However, such an involvement of K-loop towards autoinhibitory regulation of kinesin-3 motors has so far not been described. Indeed, the autoinhibitory effects observed between the K-loop of KIF1A and DNA elements of 6HB-400nm scaffold are of non-physiological nature. However, a stretch of acidic amino acids is present in the coiled-coil regions of kinesin-3 motors that can mediate such an electrostatic interaction. Additionally, the MAP mediated regulation of kinesin-3 family motors might also involve such an electrostatic interaction (Monroy et al., 2020). Dimerization of kinesin-3 motors has been proposed to be an important regulatory step for their motility (Soppina et al., 2014; Al-Bassam et al., 2003; Patel et al., 2021; Siddiqui and Straube, 2017; Hammond et al., 2009). Since multimerization of kinesin-3 monomers can also lead to intracellular cargo transport (Schimert et al., 2019), here we propose an additional tier of autoinhibitory regulation of kinesin-3 motors mediated by the K-loop. Autoinhibition of motors by the cargo domain is a common feature among kinesin family motors (Sweeney and Holzbaur, 2018). In this study using DNA origami and ensemble motors we have illuminated an autoinhibition mechanism that was not described previously. Further underscoring the power of studying motors in ensembles and the importance of the 6HB-400nm DNA origami scaffold developed in this study.

## Materials and Methods

### Protein Purification

Truncated Rat KIF1A (1-393 amino acids, dimer) followed by a GCN4 leucine zipper and Rat KIF1A (1-369 amino acids, monomer) was cloned into a pET-17b vector with a SNAP-tag followed by a 10X Histidine-tag at the carboxy-terminus. All the KIF1A gene constructs were expressed using the BL21(DE3) bacterial expression system. Transformed cells were grown at 37°C to 0.D 0.4-0.6 followed by induction with 0.5mM IPTG and overnight shaking at 18°C. Cells were harvested and lysed in Buffer-A containing 25mM pipes, pH 6.8, 100mM KCl, 5mM MgCl2, 5mM β-mercaptoethanol, 30mM imidazole. The supernatant was loaded onto a Ni-NTA column, followed by a high salt wash (Buffer-A with 300mM KCl, 200μM ATP) followed by elution using Buffer-A with 350mM imidazole. Pure proteins were obtained by further subjecting the Ni-NTA elute to size exclusion chromatography using an S200 16/1600 column (GE) with Buffer-A.

### Preparation of Benzyl Guanine-labeled oligo

Benzyl-guanine NHS ester (BG-GLA-NHS; NEB S2040) was covalently linked to the C7-amine modified oligonucleotide (Sigma), LPAH2 (see Supplementary Table 1). Briefly, 2mM LPAH2 (purchased from Sigma), was mixed with 20mM BG-NHS (dissolved in DMSO) in a molar ratio of 1:30 in the presence of 65mM HEPES pH 8.6 and incubated at 37°C with constant stirring overnight. The mixture was subjected to speed vacuum to get rid of the DMSO and further reconstituted in water. BG-labelled oligo was separated from excess BG-NHS and unlabeled oligo by subjecting this mixture to reverse phase HPLC using C18 column. Briefly, 100uL of the oligonucleotide mixture was injected into a clarity 5u oligo RP column (Phenomenex 00B-4442-E0) and was subjected to an increasing gradient of acetonitrile starting from 5% to 35% in 0.5M TEAA in 30 minutes. The NH2-oligo and the BG-labeled oligo were separated during this elution. Labeling was confirmed by mass spectrometric analysis. The BG-labeled oligonucleotide peak was collected, speed vacuumed, and further purified using obtained ethanol precipitation.

### Labelling Kinesin with Benzyl-Guanine oligonucleotide (BG-oligonucleotide)

SNAP-tagged KIF1A proteins were incubated with a 2X molar excess of BG-labeled oligonucleotide at room temperature for 15 minutes followed by overnight incubation at 4°C. The removal of excess BG-oligonucleotide from the reaction was achieved by subjecting the motor to Microtubule-binding and its release in the presence of ATP. Active BG-labeled oligonucleotide motors obtained were flash frozen as 5uL aliquots and stored at -80°C. In case of dimeric kinesin, excess BG-oligo removal was achieved by another round of Ni-NTA purification. Labeling efficiency was assessed by a 10% SDS-PAGE gel as labeled protein showed a distinct gel shift (Supplementary Figure 3).

### 6HB-400nm DNA origami structure preparation

The 6HB-400nm structure was modified from a previously published structure (Bui et al., 2010). The sequences of all staple strands, and anti-handle stands are given in the Supplementary Table 1. 10nM m13mp18 ssDNA (NEB S4040) were mixed with 100nM core staples and 500nM handle staples to a total volume of 50uL in 1X folding buffer (40mM Tris, 20mM acetic acid, 1mM EDTA, 12.5mM MgCl2, pH 8.0), followed by annealing in a PCR machine as follows: 90°C for 10 minutes, then cooled to 65°C with 1°C per minute, then further cooled to 10°C with 1°C per minute. The folded 6HB were purified from excess staples using Amicon centriprep column.

### Motility and rigor assays

Typically, a 5uL aliquot of 6HB-400nm (∼2nM) was mixed with 2uL of BG-labeled oligonucleotide kinesin (∼1uM) and incubated for 1 hour at RT and subsequently in ice for the duration of the assay. Successful formation of motor-6HB complex was assessed by 1% agarose gel where complexes showed graded retardation in mobility with increasing number of motors in the complex (Figure 1 and Supplementary Figure 3).

Biotinylated-CY5-labelled Microtubules were attached onto an acid washed glass surface using biotin-streptavidin chemistry. 0.9uL of motility mixture (motor-6HB-400nm complex in 1XBRB80 containing 1mg/mL Casein and 20μM Taxol, 2.5mM PCA, 50nM PCD, 50nM Trolox and 2mM ATP) was added to the motor-6HB-400nm complex and flowed onto the motility chamber containing microtubules. In the case of rigor experiments, ATP was omitted from the motility mixture. For rescue experiments, respective oligonucleotides as indicated was added to the motor:6HB-400nm mixture, for detailed information regarding the sequence of oligonucleotides see Supplementary Table1. Single-molecule motility of DNA-motor complex was imaged using a Nikon Ti-2 microscope (1.49 N.A., 100x objective) using total internal reflection microscopy (TIRF); images were acquired with a Hamamatsu sCMOS camera and NIS Elements software at 1-5Hz intervals. Velocities were calculated using kymograph analysis in ImageJ software and Fiesta software.

## Figure Legends

**Supplementary Figure 1:**
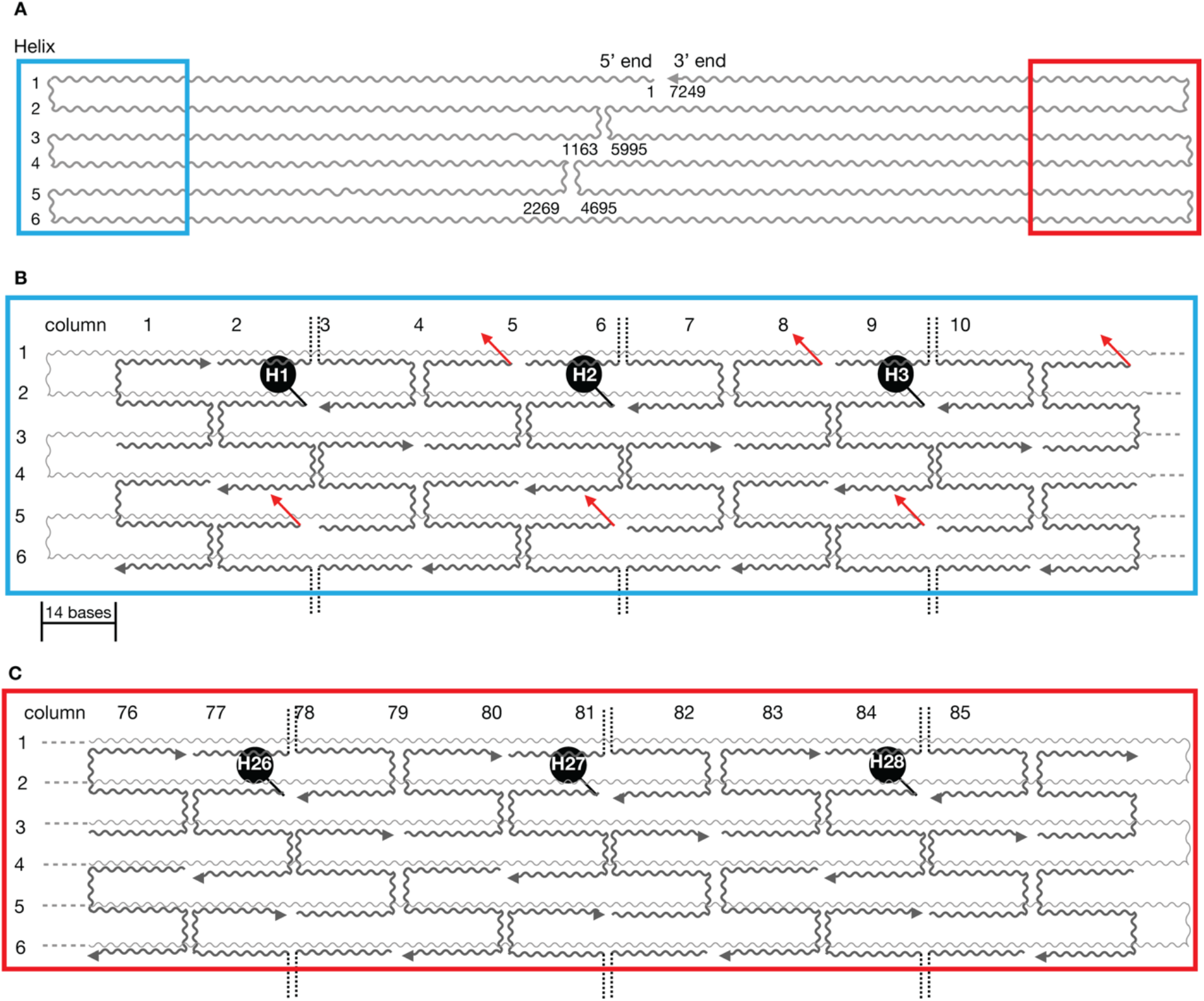
Illustration of DNA scaffold with protein attachment and fluorescent handle sites. **A**. Arrangement of m13mp18 single-stranded DNA as 6HB-400nm DNA origami scaffold. The helices and corresponding base number are as indicated. **B & C**. Zoomed in view of two distal end of 6HB-400nm DNA origami scaffold as color coded in A. The m13mp18 strand and the complementary staple oligos are marked as wavy lines in light and dark grey respectively. The fluorescent handle extensions are indicated as red arrows. The helices, column number and handle numbers (H1, H2, H3…H26, H27 & H28) and nucleotide base resolution are as indicated.

**Supplementary Figure 2:**
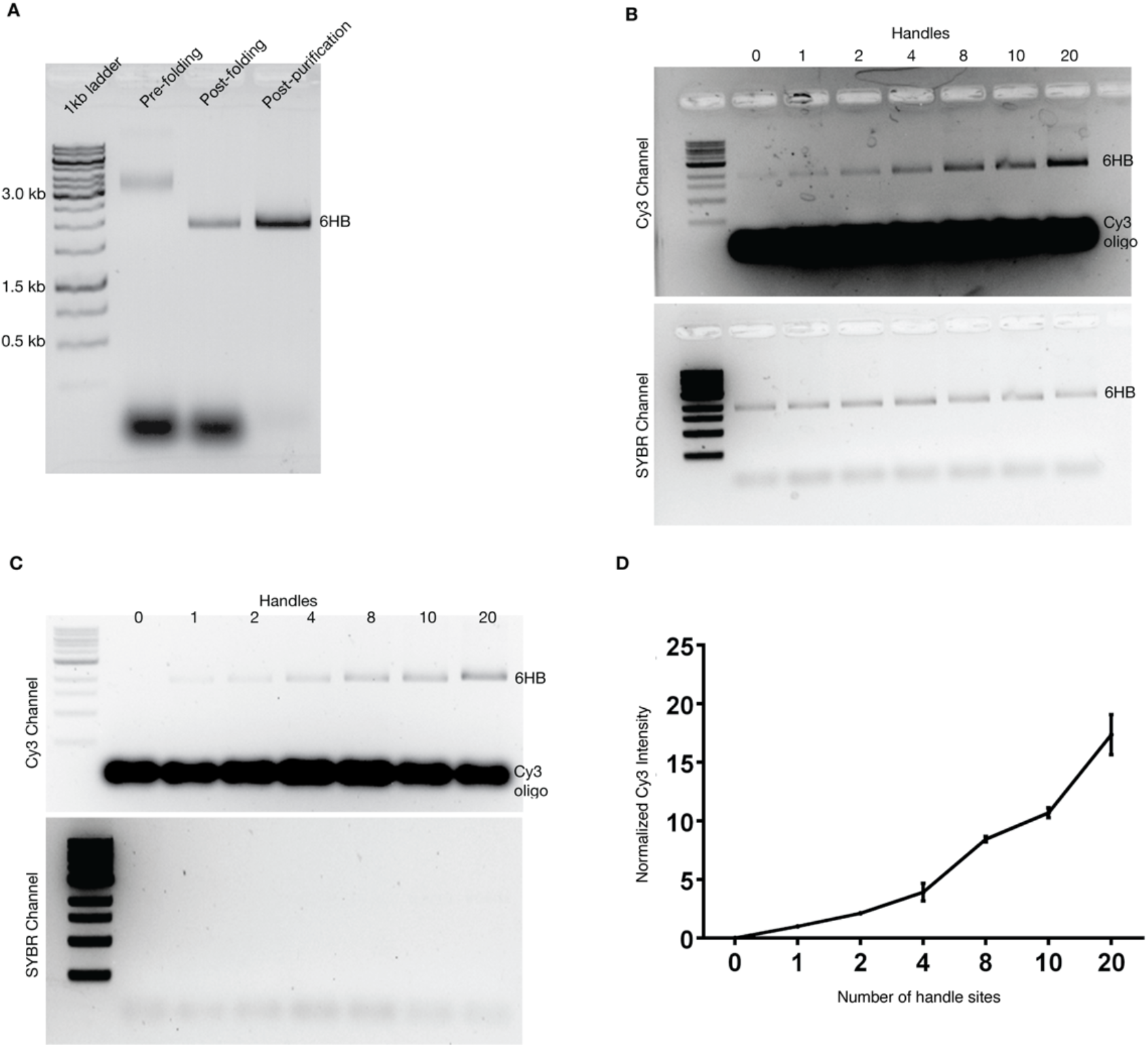
Folding and validation of 400nm-6HB handle occupancy. **A**. Agarose gel of m13mp18 scaffold (pre-folding lane), folded reaction of 400nm-6HB (post-folding lane) and 6HB-400nm after removing the excess oligonucleotides (post-purification lane). The is marked as 6hb and the size of ladder as indicated. **B**. Agarose gel of 6HB-400nm with varying handle numbers incubated Cy3-labelled anti-handle oligonucleotides as indicated, and the same gel was imaged under Cy3 and SYBR (UV) channel. **C**. Agarose gel of 6HB-400nm with varying handle numbers incubated Cy3-labelled anti-handle oligonucleotides as indicated. To rule out effects of the SYBR green, the experiments were performed without SYBR. The same gel was imaged under Cy3 and SYBR (UV) channel. **D**. Mean fold increase in Cy3 fluorescence as a function of handle number present in the 6HB-400nm DNA origami scaffold. Error bars represent the standard error of the mean from three independent experiments.

**Supplementary Figure 3:**
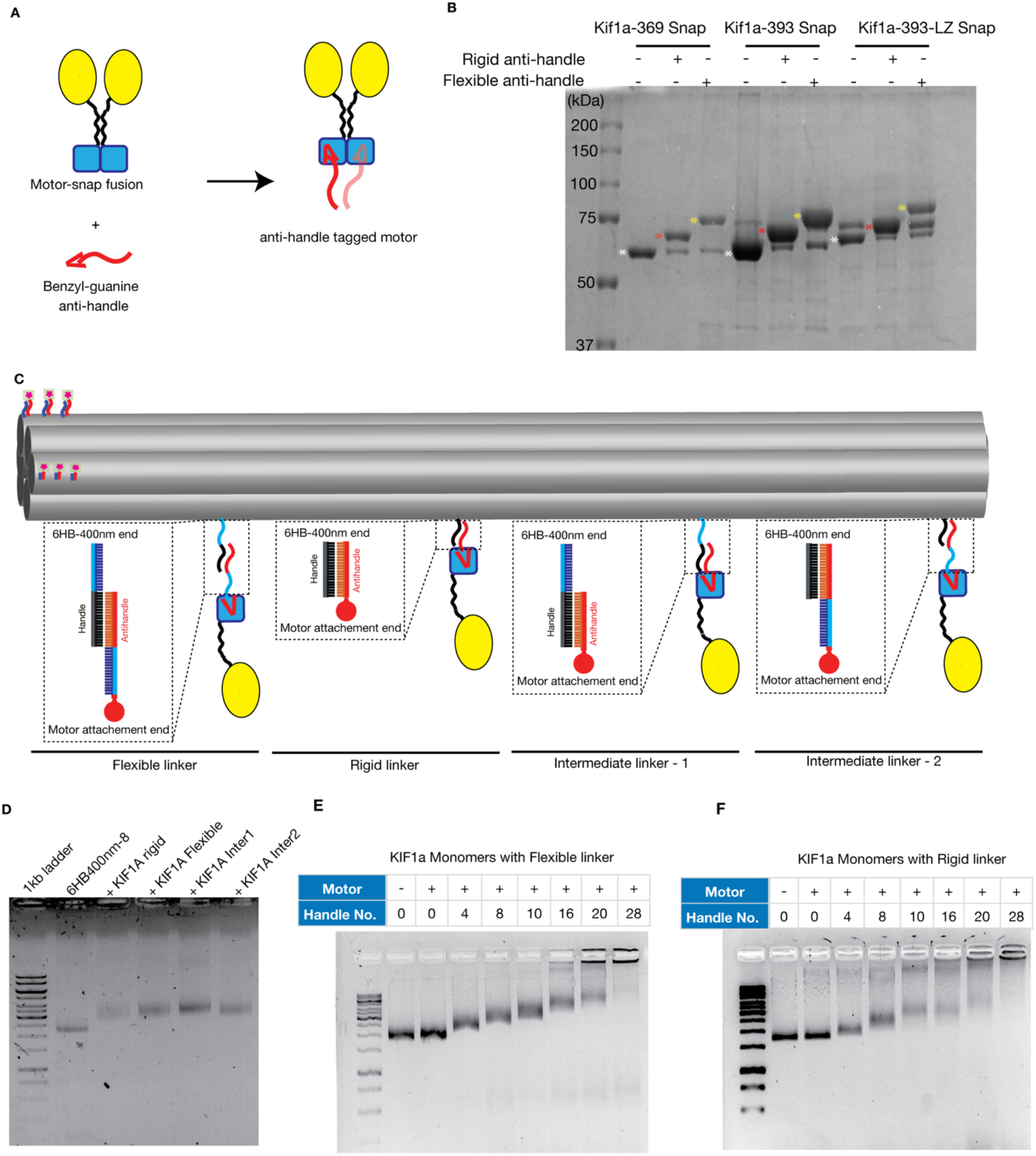
Engineering kinesins for DNA attachment. **A**. Cartoon representation of KIF1A motors tagged with oligonucleotides using benzyl-guanine and SNAP tag method. **B**. SDS-PAGE gel of KIF1A monomer and dimer before and after tagged with flexible and rigid anti-handle oligonucleotides as indicated. The white, red and yellow asterisks denote KIF1A unconjugated, KIF1A conjugated with rigid and flexible oligonucleotides respectively. **C**. Illustration of 6HB-400nm DNA origami scaffold with flexible, rigid and intermediate motor linkers as indicated. For details regarding the sequences of oligonucleotides see Supplementary Table 1. **D**. Agarose gel shift assays for 6HB-400nm scaffold with and without KIF1A monomer ensembles for the 8H used in Figure 3 rigor experiments. **E. & F**. Agarose gel shift assays for 6HB-400nm scaffold with and without KIF1A monomers with varying handle sites for flexible and rigid linkers as marked.

**Supplementary Figure 4:**
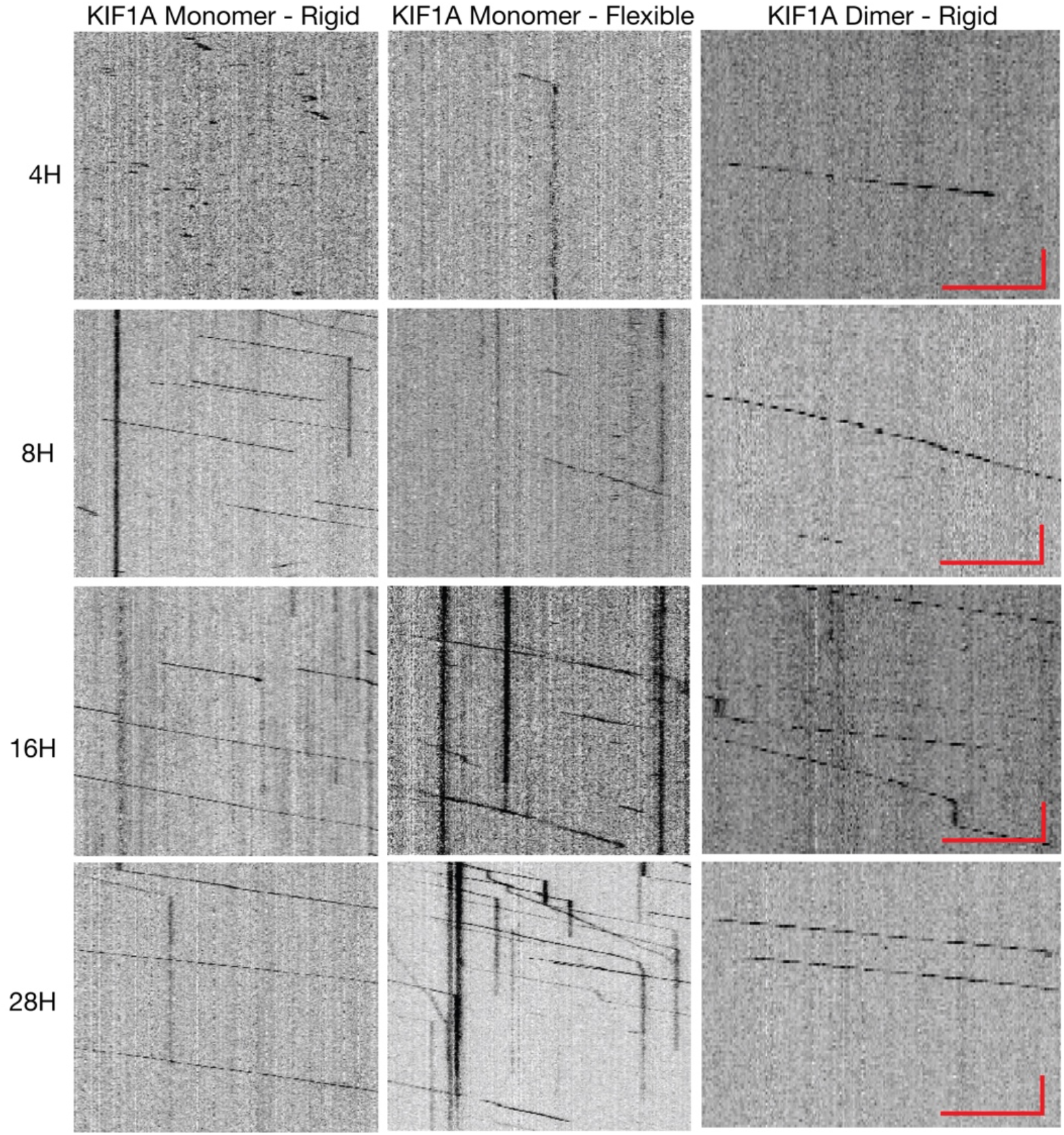
Kymographs of KIF1A ensemble motility. Representative kymographs of KIF1A ensembles with flexible and rigid anti-handle oligonucleotides, for the data shown in Figure 2 as indicated. Scale bar = 5μm and 10 seconds.

**Supplementary Figure 5:**
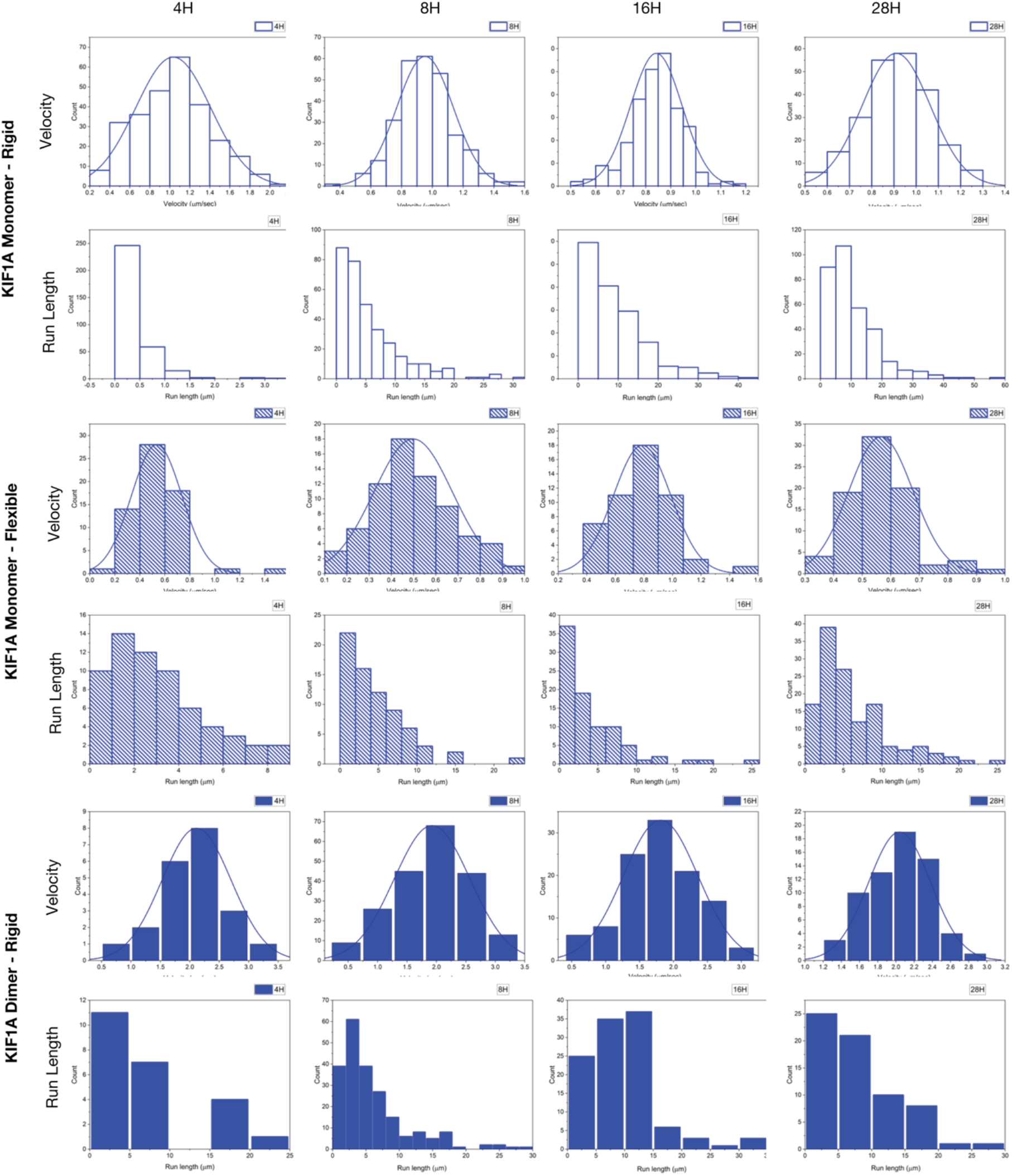
Velocity and run length histograms of KIF1A ensemble motility. Velocity and processivity histograms of 6HB-400nm 4-28H KIF1A monomer and dimer ensembles. Data for KIF1A monomer ensembles with flexible, rigid and KIF1A dimer ensembles with rigid anti-handle oligonucleotides are shown as indicated.

**Supplementary Figure 6:**
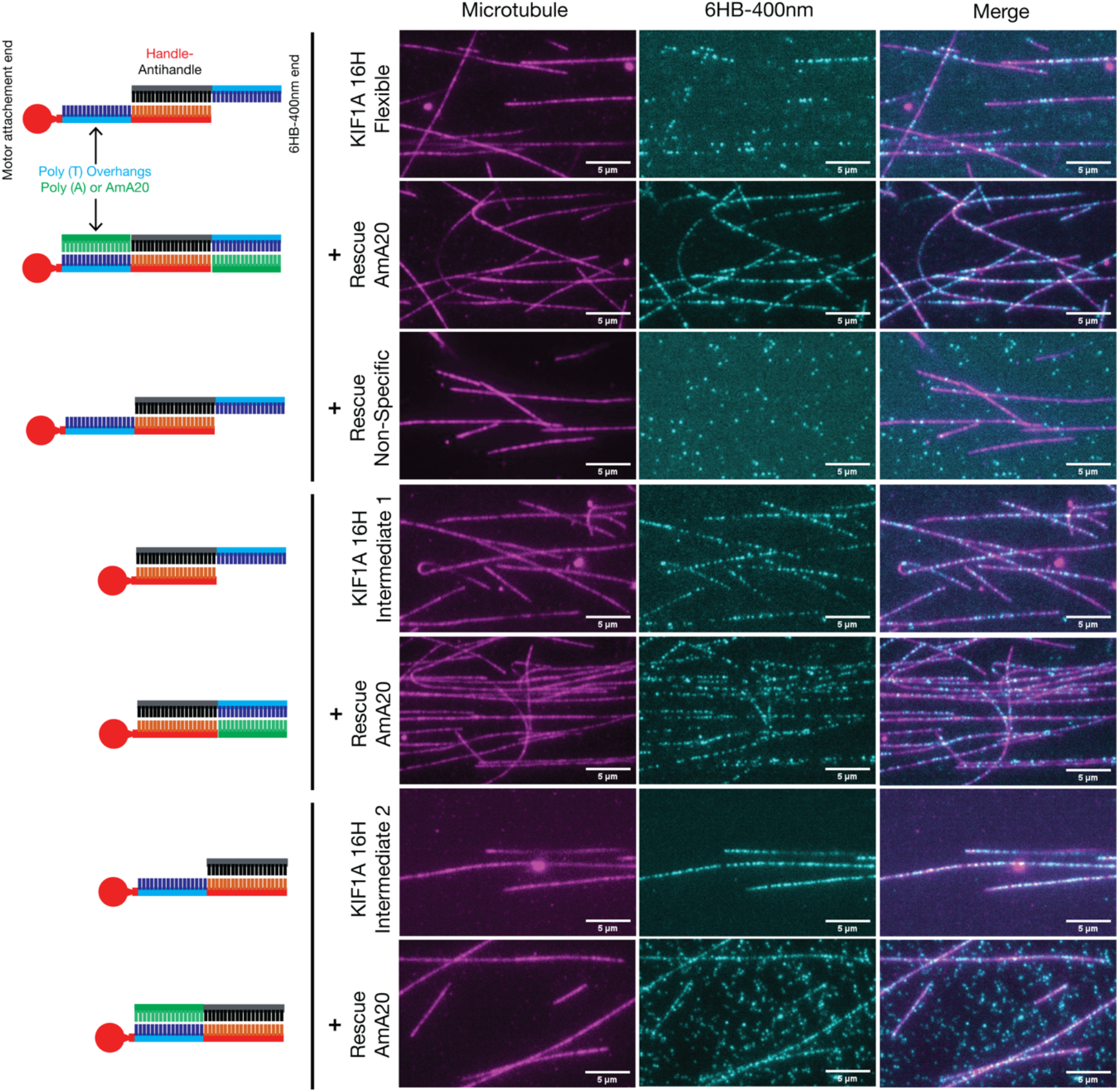
Microtubule binding of KIF1A 16H ensembles. Microtubule-binding of 6HB-400nm 16H KIF1A monomer ensembles with varying linkers as indicated and illustrated. The microtubules are shown in magenta and the 6HB-400nm KIF1A ensembles in cyan. In each microtubule-binding experiment, a rescue reaction was perfumed (marked as rescue), where an oligonucleotide that is complimentary to the flexible parts of linkers was added to the mixture. For more details regarding the sequences of linkers and rescue oligonucleotides see Supplementary Table 1. Scale bar = 5μm.

**Supplementary Figure 7:**
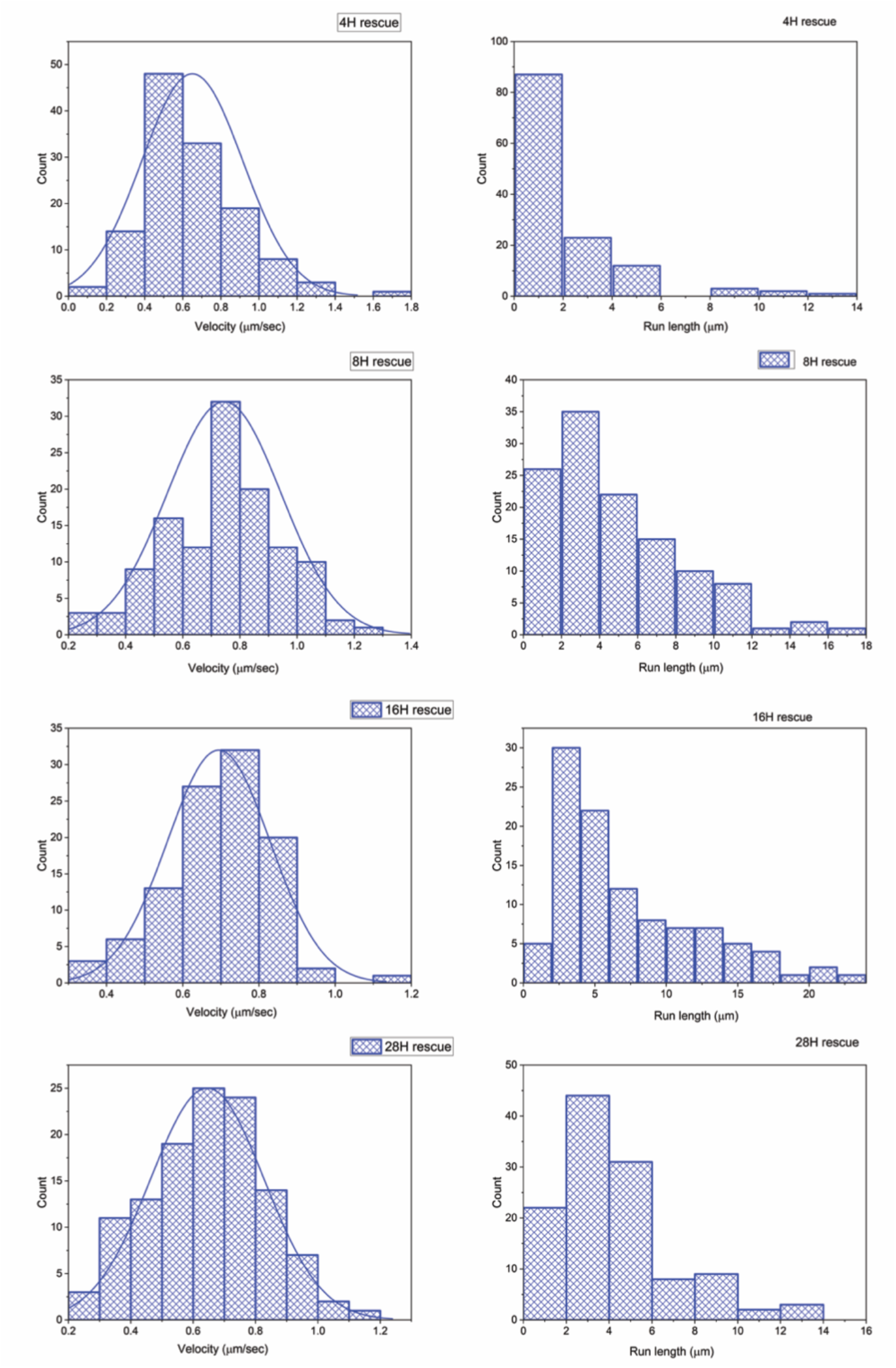
Histograms and kymographs of KIF1A ensembles with flexible and rescue linkers. **A**. Velocity and processivity histograms of 6HB-400nm 4-28H KIF1A monomer ensembles with flexible and flexible-rescue oligonucleotides, for the data shown in Figure 4.

**Supplementary Movie 1:** KIF1A 8H ensembles with rigid, flexible and flexible with rescue oligonucleotides as indicated. Scale bar = 5μm and time in minutes/seconds (mm:ss format) as indicated.

## Acknowledgments

We thank the Sirajuddin lab members for critical feedback on the manuscript. P.L acknowledges the support from inStem Graduate Program. M.S acknowledges funding support from inStem core grants from the Department of Biotechnology, India, a DBT/Wellcome Trust India Alliance Intermediate Fellowship (IA/I/14/2/501533), EMBO Young Investigator Programme award, CEFIPRA (5703-1), the Department of Science and Technology, SERB-EMR grant (CRG/2019/003246) and DBT-BIRAC (BT/PR40389/COT/142/6/2020).

## Author Contribution

P.L and M.S designed the work. P.L performed the experiments and analyzed the data. M.S supervised the work and wrote the paper.

